# Thermodynamic modeling reveals widespread multivalent binding by RNA-binding proteins

**DOI:** 10.1101/2021.01.30.428941

**Authors:** Salma Sohrabi-Jahromi, Johannes Söding

## Abstract

**Motivation:** Understanding how proteins recognize their RNA targets is essential to elucidate regulatory processes in the cell. Many RNA-binding proteins (RBPs) form complexes or have multiple domains that allow them to bind to RNA in a multivalent, cooperative manner. They can thereby achieve higher specificity and affinity than proteins with a single RNA-binding domain. However, current approaches to de-novo discovery of RNA binding motifs do not take multivalent binding into account.

**Results:** We present Bipartite Motif Finder (BMF), which is based on a thermodynamic model of RBPs with two cooperatively binding RNA-binding domains. We show that bivalent binding is a common strategy among RBPs, yielding higher affinity and sequence specificity. We furthermore illustrate that the spatial geometry between the binding sites can be learned from bound RNA sequences. These discovered bipartite motifs are consistent with previously known motifs and binding behaviors. Our results demonstrate the importance of multivalent binding for RNA-binding proteins and highlight the value of bipartite motif models in representing the multivalency of protein-RNA interactions.

**Availability:** BMF source code is available at https://github.com/soedinglab/bipartite_motif_finder under a GPL license. The BMF web server is accessible at https://bmf.soedinglab.org.

## Introduction

RNA molecules in the cell are rarely naked but rather covered with numerous RNA-binding proteins (RBPs) (35). These RBPs play a crucial role in regulating the various steps of RNA biochemistry, from RNA maturation and transport, to cellular localization, translation, and degradation (7). RNA molecules can in turn regulate RBP function by altering their stability, interaction partners, and localization (12). These processes require specific binding of RBPs to their target RNAs. RBPs mostly achieve this specificity through RNA-binding domains (RBDs) that engage with specific RNA sequences and\or structures (20). Unraveling the target preferences of RBPs is therefore key to understanding cellular regulation.

Many experimental techniques have emerged to generate systematic maps of protein-RNA interactions. To find *in-vivo* binding sites, many variants of RNA immunoprecipitation (RIP-seq) (9) and cross-linking immunoprecipitation (CLIP-seq), such as PAR-CLIP (10), iCLIP (18) and eCLIP (39), have been proposed. In both approaches, RNAs bound to the immunoprecipitated protein of interest are sequenced and mapped to the genome. Deriving accurate models of binding affinities from *in-vivo* data is problematic because RBP-RNA interactions are influenced by cooperativity and competition with other RBPs, and are additionally influenced by RNA localization, expression, and folding (2). Therefore, techniques have been developed to measure binding affinities *in-vitro*, in isolation from other RBPs, using random libraries of RNA substrates: RNA Bind-n-Seq (RBNS) (19), RNA-compete (30, 5), and high-throughput RNA-SELEX (HTR-SELEX) (13).

A wide range of motif discovery tools have been developed to learn models of sequence- and secondary structure-dependent binding affinties of RBPs based on datasets of sequences bound in-vitro or in-vivo by an RBP of interest (15, 23, 38, 25). More recently, a new wave of algorithms have been introduced that use deep neural networks to predict RBP binding sites (1, 3, 28, 8). While these deep learning methods have promising accuracy in predicting RBP binding, interpreting what these networks have learned remains challenging. Moreover, with the increasing number of model parameters and network complexity, the risk grows that such models could also learn experimental biases in the datasets. This is particularly problematic for RBPs, since many of them show short and degenerate sequence preferences. Moreover, RBPs often bind low-complexity untranslated regions in the RNA (6), unlike transcription factors, which usually bind to more complex sequence motifs and have higher binding specificities. RBPs have further been shown by spaced k-mer counting approaches to often bind with multiple RNA-binding domains two separated cores with usually similar or identical motifs (6,13). A recent deep learning software is the only available one capable of learning distance dependent motif pairs (29).

In this work, we present Bipartite Motif Finder (BMF), a tool for learning bipartite RNA motifs in RNA-ptotein interaction datasets. To accurately model the binding affinity of RBPs possessing domains with similar core motifs of low-information content that bind to RNAs with a relatively high density of the core motifs, BMF sums up the contribution of all alternative binding conformations, and not just the best binding configuration. To the best of our knowledge, BMF is the first approach that adopts a thermodynamic viewpoint to RBP *de novo* binding motif discovery. We demonstrate that BMF is able to detect short and degenerate motifs and to learn the spatial relationship between them. We furthermore show that around half of RBPs manifest multivalent binding with a preferential linker distance between the two binding sites.

Benchmarking the performance of learned binding site models by cross-validation can be problematic when testing methods that train highly parameterized models such as deep neural networks, as these methods can learn biologically irrelevant sequence biases inherent to the experimental method. To compare BMF to existing tools and assess their capacity for learning relevant motif sequences that predict binding events in the cell, we built a cross-platform validation benchmark, training models on HTR-SELEX data and testing on *in-vivo* CLIP data. Despite the many complicating effects in vivo, we find that the motif and distance preferences learned by BMF can predict RBP binding in the cellular context and that high-quality motifs learned in *vitro* are often very similar to the motifs learned on *in-vivo* data. Moreover, BMF can predict binding sites on par with or even better than existing tools.

## Methods

Most RBPs can bind RNA using several structured RBDs and often also using disordered regions, some of which contain typical RGG/RG and RS motifs, which can modulate RNA-binding activity (21, 4, 27). Furthermore, many RNA-binding proteins dimerize or homo- and hetero-oligormerize. This effectively leads to two and more RNA-binding domains binding cooperatively to RNA molecules. Here, we present Bipartite Motif Finder (BMF), a motif search tool and algorithm to describe the sequence specificity of monovalently and multivalently binding proteins or protein complexes.

### Thermodynamic model for bivalent RNA binding

We consider the simple case in which the RBP consists of two RBDs, *A* and *B* (Figure 1A). We describe the binding of proteins at concentration *c*_AB_ to a single, specific RNA sequence **x** = (*x*_0_… *x*_*L*–1_) = *x*_0:*L*–1_ composed of nucleotides *x_i_*. We consider not only the most likely binding configuration but rather all possible binding configurations, involving zero, one or more proteins bound to the RNA (Figure 1B). According to Boltzmann’s law, each binding configuration **c** has a probability *p*(**c**) proportional to its so-called *statistical weight e*^(−*E*(**c**)−*T*Δ*S*(**c**))/*k*_B_*T*^, where *F*(**c**) = –*E*(**c**) – *T*Δ*S*(**c**) is the free energy composed of the binding enthalpy –*E*(**c**) and a part related to the change in entropy Δ*S*(**c**) between the completely unbound and bound states. To obtain probabilities, the statistical weights need to be normalized at the end by dividing by their total sum, the partition sum *Z*(**x**).

**Figure 1:**
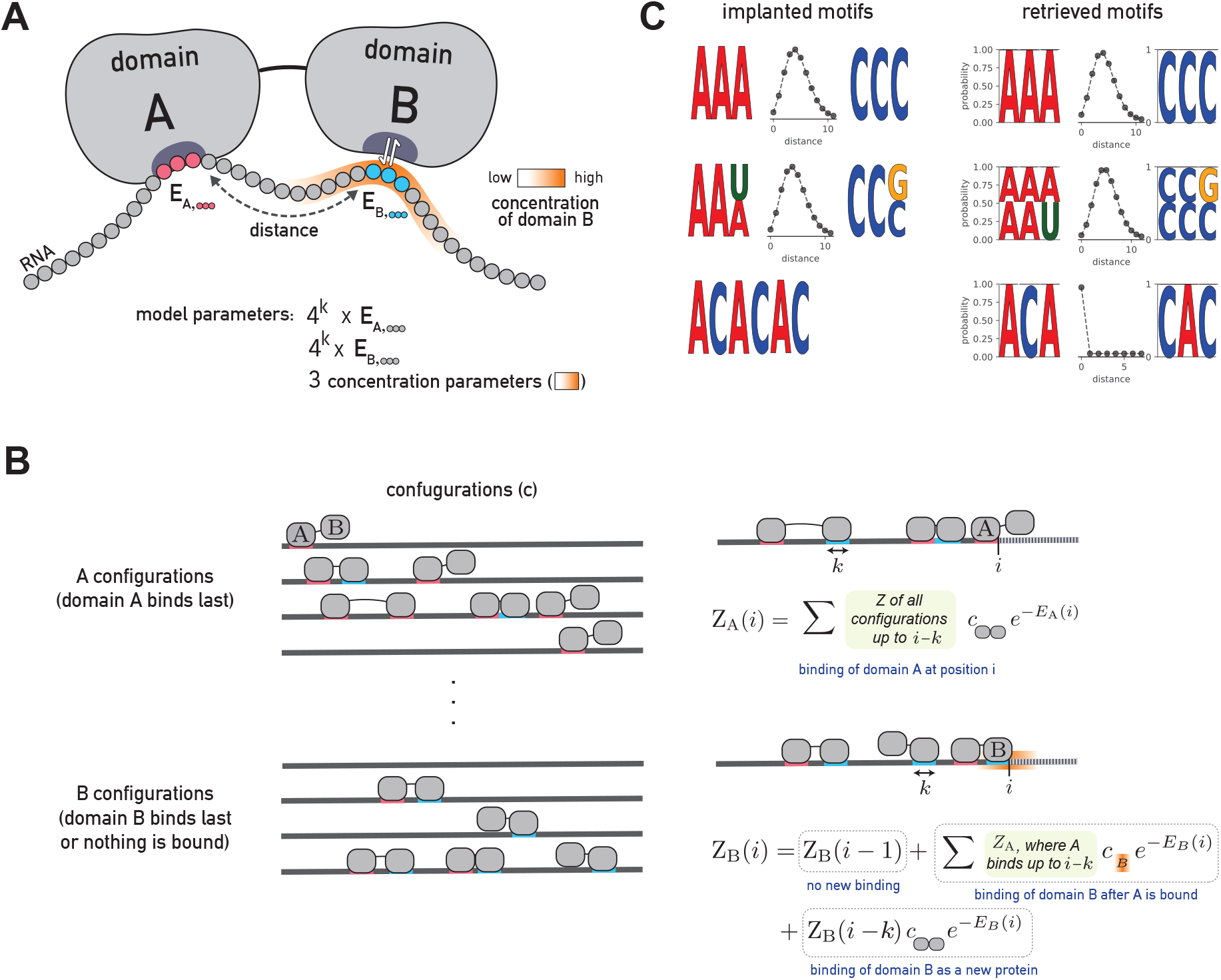
BMF can learn multivalent binding preferences for RBPs. **(A)** RBP-RNA interaction model for a protein with two RNA-binding domains. BMF optimizes the binding energies of each domain to all possible RNA *k*-mers (*k* = 3 here) and learns the distance distribution between the motif cores. BMF models the high RNA local concentration at the second binding site, when the first domain is bound to the RNA. **(B)** BMF calculates binding probabilities for all binding configurations of one or several proteins to the RNA sequence. *Z*_A_(*i*) is the sum of statistical weights of all binding configurations on the RNA up to position i, for which domain A is bound at position i. Similarly, *Z*_B_(*i*) is the sum of statistical weights of all binding configurations on the RNA subsequence for which no domain is bound or domain B is bound with its right edge upstream of or at position *i*. *Z*_A_ and *Z*_B_ are calculated iteratively (right panel). The first term in the second equation accounts for configurations for which position *i* is not bound by anything, the second term accounts for configurations for which domain A of the same protein is bound at *j* (as seen in the example illustration) and the last term accounts for configurations for which domain B binds whose A domain is not bound upstream of *i*. **(C)** BMF recovers the correct RNA motifs implanted in synthetic datasets for all tested cases. Here and in the following figures, the two learned core motifs are visualized by plotting the energies of the top five k-mers, converted to *k*-mer probabilities according to Boltzmann’s law and normalized to 1.

The change in entropy due to the binding of a single protein that is present at concentration *c*_AB_ is equal to its chemical potential, which is Δ*S* = *k*_B_ log *c*_AB_. In the following, we compute all energies in units of *k*_B_*T*, so we set *k*_B_*T* = 1. In our model, the concentration *c*_B_(*d*) of the downstream domain *B* at the RNA depends on the distance *d* to the binding site of the upstream domain *A* (see next subsection).

We compute the statistical weights of all binding configurations iteratively using dynamic programming. We split the configurations into two sets, A and B, and define *Z*_A_(*i*) to be the sum of statistical weights of all binding configurations on the RNA up to position i, *x*_0:*i*_, for which domain A is bound at position *i* – *k* + 1 to *i*, where *k* is the length of RNA bound by the domains. Similarly, we define *Z*_B_(*i*) to be the sum of statistical weights of all binding configurations on the RNA sequence *x*_0:*i*_ for which no domain is bound or domain B is bound with its right edge upstream of or at position *i*. With the knowledge of *Z*_A_(*i′*) and *Z*_B_ (*i′*) for 0 ≤ *i′* < *i*, we can compute *Z*_A_(*i*) and *Z*_B_(*i*) (Figure 1B):

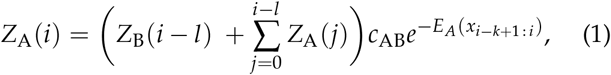

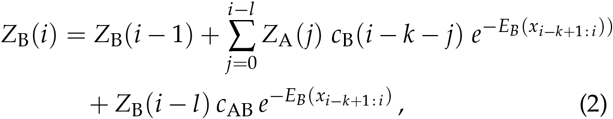

where *E_A_*(*x*_*i*–*k*+1: *i*_) and *E_B_* (*x*_*i*–*k*+1: *i*_) represent the binding energies of domains A and B to the RNA sequence *x*_*i*–*k*+1: *i*_. The concentration of the single B domain, defined as expected number of B per volume, is simply its probability density. The dynamic programming is initialized using

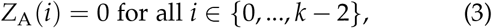

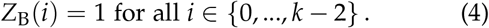

The first equation follows from requiring all *k* positions in the binding motif to be part of sequence *x*_0:*L*–1_. The second equation follows from the fact that *Z*_B_(*i*) for *i* < *k* – 1 sums up only the statistical weight of the unbound configuration.

The partition sum *Z*(**x**) for RNA sequence **x** is the sum of statistical weights of all configurations,

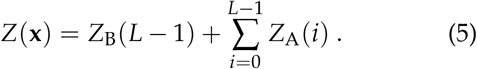

The probability for an RNA to not be bound by any protein (neither *A* nor *B* domains) is just the statistical weight of the unbound configuration, set to 1, times the normalization factor 1/*Z*(**x**), so the probability for a RNA **x** to be bound by a protein is *p* (bound |**x**) = 1 – 1/*Z*(**x**).

By taking the partial derivatives of equations (1) and (2) with respect to the model parameters (Supplementary Methods), we obtain update equations for the partial derivatives with which we can in turn compute the partial derivatives of *Z*(**x**), *p*(bound|**x**), and the log likelihood in equation (8). These allow us to find optimum model parameters by gradient-based maximization of the log-likelihood.

### Motif model of a single RNA-binding region

Position weight matrices (PWMs) and Bayesian Markov models (BaMMs) have been used to represent RBP binding preferences through positional or conditional probabilities of observing each nucleotide at a given position (11, 34). Since RBPs are known to bind shorter and more repetitive sequences, we learn binding energies for all 4^*k*^ *k*-mers at each motif core, *E_A_* (*k*-mer) and *E_B_* (*k*-mer). The length *k* of the motif can be set by the user.

### Model for the effective concentration *c*_B_ (*d*)

Spaced *k*-mer analyses on high-throughput RNA-binding datasets pointed to a length preference of the RNA linker connecting two motif cores (13, 6, 32). The concentration of domain *B* after domain *A* binds the RNA molecule is equal to its probability distribution. While according to the flexible chain model of the RNA fragment the concentration should be a Gaussian distribution centered on domain A (31), for short RNA linkers the concentration can peak some distance away from domain A. To describe multivalent binding for both short-range and long-range co-occurrence of motif sequences, we model the effective concentration at the second binding site with a negative binomial (NB) distribution,

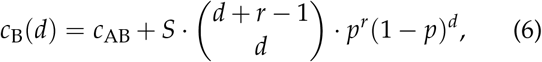

where *d* represents the the number of nucleotides between the binding sites of A and B on the RNA, and *r* and *p* are parameters of the negative binomial distribution. The total concentration of *B* is the cellular concentration (*c*_AB_) plus *c*_B_(*d*), the local concentration of *B* linked to a bound *A*. We scale the negative binomial with the factor *S* as a conversion to protein concentration values. Since only the ratio between *S* and *c*_AB_ determine the binding dynamics, we fix *c*_AB_ to one and optimize our bipartite model for *S, r*, and *p*.

### Parameter initialization

The absolute values of the energy parameters in our model do not reflect the physical binding energies, however their relative values determine the probability of binding to a given sequence. We therefore draw initial energy parameters randomly (in units of *k*_B_*T*) from a normal distribution with the average of 12 and standard deviation of one. The initial value of 12 kB *T* was chosen based on experimentally determined binding energies (43) and additionally ensures that the algorithm does not overflow. The scaling factor *S* is initialized as 10^4^, The spacer parameter *r* is drawn from a uniform distribution from one to five and *p* is randomly drawn between zero and 0.5.

### Likelihood estimation for HTR-SELEX measurements

In HTR-SELEX experiments (and similarly for bind-n-Seq), we have input (background) library sequences 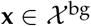 and sequences enriched after competitive binding, 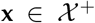. We denote with *p*_b_(**x**) the fraction of sequence **x** in the input library. To find a sequence in 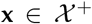, it must have first been present in the input library (probability *p*_b_(**x**)) and then have been bound to the RNA (probability *p* (bound |**x**)). The probability to find a sequence 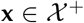 after the selection is therefore, according to Bayes’ theorem,

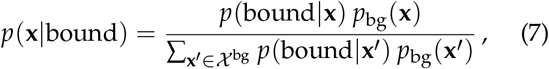

and, using *p*(bound |**x**) = 1 – 1/*Z*(**x**), the log-likelihood is

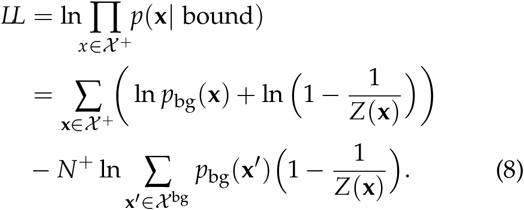

### Parameter optimization

We learn the model parameters by maximizing the likelihood function (eq. 8). For an efficient optimization using stochastic gradient descent, we computed the partial derivative of the likelihood function with respect to all of the model parameters (Supplementary Methods). For parameter optimization, we used ADAM (16) with hyperparameters *α* = 0.01, *β*_1_ = 0.9, *β*_2_ = 0.999, and *ε* = 10^−8^, and a minibatches size of 512. We parameterized *r* = *e^ρ^* and *p* = 1/(1 + *e^−π^*) to ensure that *r* and *p* stay within bounds. Optimization was terminated when 1000 iterations were reached or when the variation *v_θ_* for the best bound *k*-mer of each domain as well as for *p* and *r* fall under a threshold of 0.03. The variation for the parameter *θ* up to iteration *t* was defined as *v_θ_* = (max{*θ*_*t*-4: *t*_} – min{*θ*_*t*-4: *t*_})/*θ_t_*.

### Evaluating the performance of BMF on synthetic data

In order to evaluate BMF’s ability to learn bipartite motifs, we generated two sets of 2000 RNA sequences, an artificial input set and an enriched set. For the enriched set, we inserted the first core of the simulated bipartite motif at random positions. The second core was inserted with a linker length drawn from a binomial distribution with a specific *p* and *r*. We ran BMF 10 times with random parameter initializations to assess its robustness.

### HTR-SELEX datasets

We obtained 177 HTR-SELEX datasets of 86 distinct factors from (13). We used sequences of the input library and the last cycle to train BMF. Even though our model describes one cycle of selection, the retrieved motifs were more prominent in the later cycles. Moreover, the cross-platform validation discussed below resulted in slightly better performance for all the tools when choosing the input and last cycles for motif detection in comparison to second and third cycles. Whenever several experimental or technical replicates were available, we built a separate model for each replicate and averaged the corresponding metric over all replicates of an RBP at the end. We used BMF’s default hyper-parameters throughout the manuscript.

### Cross-platform validation of in-vitro motifs

Each experimental technique for measuring RNA binding has its own biases. When measuring the quality of predictions of motif models by cross-validation, methods can learn these biases to distinguish bound from background sequences. Highly parameterized models could learn such subtle, complex biases. These platform-dependent biases can be a result of library preparation, amplification, or can depend on the type and concentration of RNase that is used (17, 26). There have been efforts to reduce the effect of such biases when training motif models, e.g. by learning binding models for many RBPs at the same time (8). In order to ensure that BMF does not over-train on the *in-vitro* HTR-SELEX data, we performed cross-platform validation: We trained BMF on HTR-SELEX datasets and used the resulting models to predict binding sites in *in-vivo* CLIP data.

We collected eCLIP datasets of 15 RBPs (40) and PARCLIP datasets of 10 RBPs (24) for which we also have HTR-SELEX data. We used the pre-processed CLIP peaks as enriched sequences. Since the PAR-CLIP dataset contained larger numbers of peaks, we restricted our analysis to the top 2000 reported binding sites per RBP. For each eCLIP and PAR-CLIP dataset, we created a background set of the same size by drawing random PAR-CLIP or eCLIP peaks of other factors measured with the same technique. We applied a sliding window with length of 50 and a stride of 20 to generate same-size fragments that fully cover each peak. The prediction scores were averaged over these fragments when the region was longer than 50 bases. We compared our simple model with deep learning approaches, the popular RBP binding predictors iDeepE (28) and GraphProt (23). iDeepE uses deep learning to predict RBP binding, while GraphProt’s model is based on Support Vector Machines (SVMs).

### BMF software and web server

The BMF command-line tool offers three commands: (1) Learning a BMF model given enriched and background sequences. Output is a BMF model file. (2) Bipartite motif visualization, given the BMF file learned in step 1. (3) Predicting binding scores for new sequences with the BMF model trained in the first step. The first two functionalities (*de novo* motif discovery) are also available on the BMF web server.

## Results

We present BMF, a method for de-novo discovery of RNA-binding motifs that uses a bipartite motif model capable of learning multivalent binding specificities among RBPs. BMF models the protein binding with up to two domains to its RNA substrate. We assume that due to the structure of the RBP (or RBP complex), the distance between the two binding sites is spatially constrained. BMF therefore consists of two short sequence motif models and a distance probability distribution (Figure 1A). Binding with just one domain is modeled using a distance distribution peaked at 0 base pairs. In the following sections we demonstrate that this model can reliably detect bipartite motifs in synthetic and real sequences, and we evaluate its performance at identifying binding sites compared to other models of RBP binding in HTR-SELEX, PAR-CLIP and eCLIP datasets.

### BMF accurately discovers implanted synthetic motifs

To test BMF’s ability to learn bipartite motifs, we generated 2000 artificial sequences containing first an AAA and then a CCC with a distance distribution of around 3 to 5 bases between them (Figure 1B, top). BMF retrieved the implanted motifs and spacer distribution accurately. The results were similarly accurate when sequence degeneracy was introduced by flipping the last base (Figure 1B, middle), or when implanting the repeat sequence ACACAC (Figure 1B, bottom). This demonstrates that BMF can not only reveal multivalent specificities but can also recover longer sequence motifs by placing the two cores adjacent to one another.

The log-likelihood increases during stochastic gradient descent and the optimization terminates when the log-likelihood has reached a plateau (Figure S1A). The distance parameters and the binding energies of *k*-mers in the motif cores all reach a plateau before termination (Figure S1B,C). To test robustness to parameter initialization, we ran BMF ten times with random initial parameter values and verified that the *k*-mer energies and distance parameters match across all runs. (Figure S1D,E).

### Most RBPs show multivalent binding, often to multiple occurrences of the same motif

We applied BMF to 177 HTR-SELEX datasets consisting of 86 distinct RBPs to investigate the importance of multivalent binding in the formation of RBP target specificity. BMF detected bipartite binding for many RBPs including ELAVL1, KHDRBS3, and RBPMS (Figure 2A). Interestingly, BMF restricted the distance of the motif cores strictly to zero when the RBP binds repeat sequences (e.g. CELF1 binding GU repeats) or when the RBP binds a longer RNA sequence that requires a longer motif core (e.g. RBFOX3 binding UGCAUG, and PUM2 binindg UGUANA). The sequence and spacing preferences were also reproducible across experimental replicates (Figure S2), and match for proteins that belong in the same family (Figure S3). All 177 BMF models with core lengths of 3-5 can be found at BMF’s GitHub repository. These results show that BMF can identify bipartite motifs in HTR-SELEX data.

**Figure 2:**
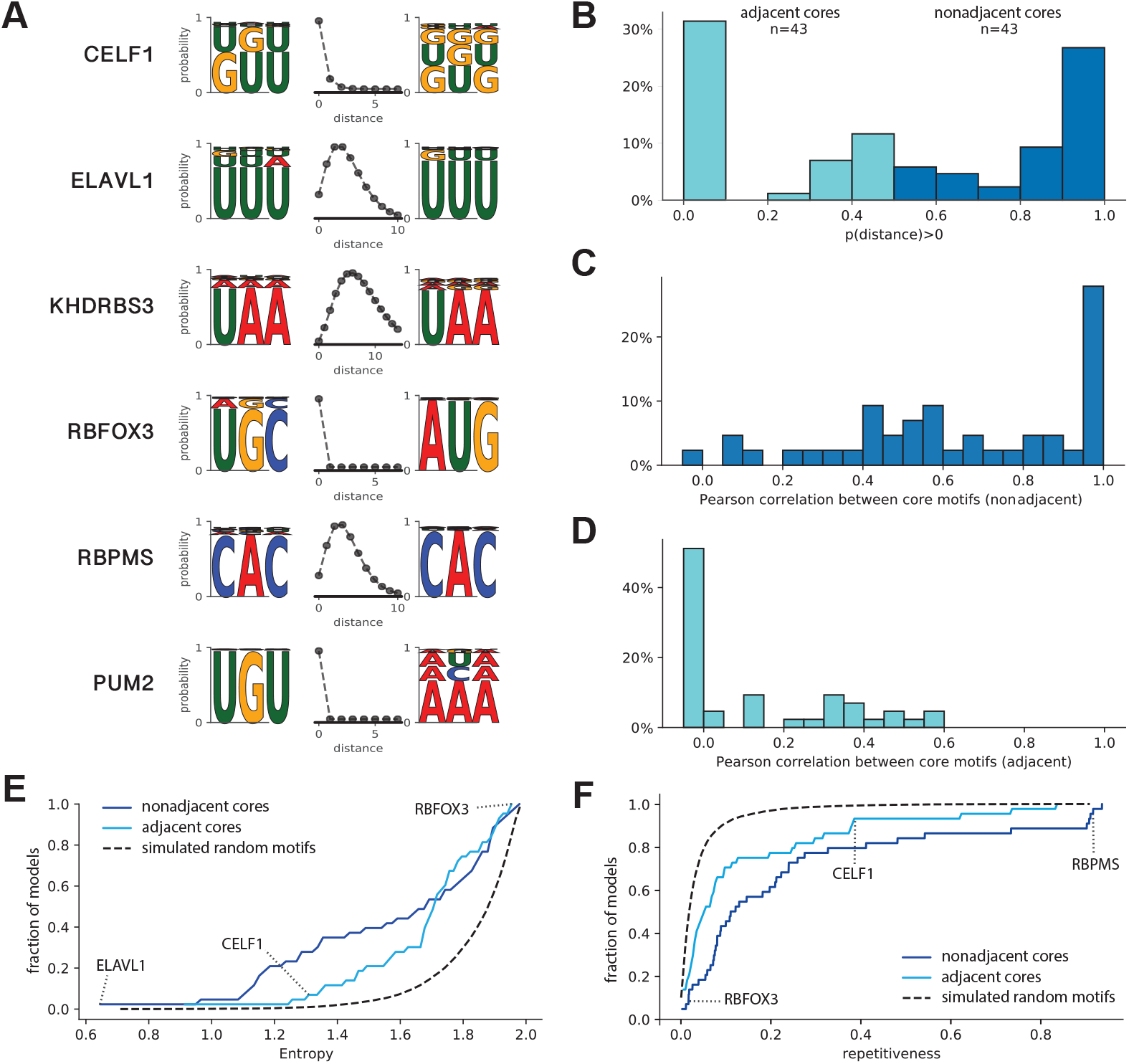
Many RBPs are multivalent, bind low-complexity sequences and often bind two similar motif cores. **(A)** Examples of motifs that represent a wide range of binding modes, learned by BMF on HTR-SELEX data. When the RBP has a larger motif than allowed by the core size (3 here), the distance between cores is learned to be zero to accommodate a longer binding sequence (e.g. CELF1, RBFOX3, and PUM2). **(B)** Distribution of the probability of the spacer length between the two motif cores to be above 0. As seen in the examples in A, most RBPs either clearly bind adjacent cores (distance= 0, turquoise) or have a multivalent binding mode with two nonadjacent cores (dark blue). **(C)** and **(D)** Similarities between binding preferences of the two cores for RBPs with adjacent cores (turquoise) or multivalent nonadjacent cores (dark blue), according to panel B. **(E)** Cumulative distribution of the entropy of BMF models for all RBPs in the HTR-SELEX dataset. In general the optimized bipartite motif models have much lower complexity than randomly generated bipartite models (dashed black line). **(F)** Cumulative distribution of *“sequence repetitiveness”* of BMF models for all RBPs in the HTR-SELEX dataset. Overall, BMF models are more often repetitive that those of randomly generated bipartite models (dashed black line).

We then looked for the frequency of such multivalent, bipartite motifs and calculated the probability of observing the two core motifs at distances beyond zero for each motif model (Figure 2B). At two extremes, this probability would be zero for RBPs like RBFOX3, which consist of a larger binding sequence, and one for RBPs like KHDRBS3, which prefer a larger spacer between the motif cores. Interestingly, the majority of RBPs lie at the two extremes, and about half of them show a bipartite binding behavior. This ratio is higher than estimated in previous studies, which were based on *k*-mer counting approaches (13, 6). The number of bipartite motifs could be furthermore underestimated as some RBPs show bipartite binding only when BMF’s core size is increased to four or five nucleotides (Figure S4). Overall, these results highlight the importance of multivalent binding as a common strategy to achieve high specificity despite having individually small and weak binding sites.

We noted that many motif models (like ELAVL1 and KHDRBS3) have similar sequence preferences on both cores. We quantified their similarity by the Pearson correlation between the probabilities of observing each of the 4^*k*^ *k*-mers. As expected from the individual examples, the core motifs are mostly similar for RBPs that exhibit bipartite binding (Figure 2C) as opposed to adjacent motif cores (Figure 2D). This demonstrates that RBPs have often evolved to bind multiple occurrences of the same or similar short sequence motifs, either using multiple same-chain RNA-binding domains or by homodimerization and oligomerization.

### RBPs often bind low-complexity and repetitive sequences

It has been shown that RBPs bind sequences of lower complexity than DNA-binding transcription factors (6, 36). This can be seen at its extreme for some of our binding models, which are composed of only one to two types of nucleotides (Figure 2A). Looking at all 78 RBP binding models, we observed that many proteins bind repetitive sequences or have the same simple *k*-mer affinities for each of their valencies. In order to quantify this, we calculated the entropy of the motif sequences as a measure of sequence complexity (Figure 2E, Supplementary Methods)(6). For highly complex sequence affinities (e.g. RBFOX3), the entropy gets close to two, while this value is closer to zero for degenerate and repetitive sequences (e.g. ELAVL1). A similar trend is visible when quantifying the repetitiveness of BMF models, resulting in high scores when both cores consist of mono- or di-nucleotide repeats (Figure 2F, Supplementary Methods). Overall, more than half of RBP motifs show levels of degeneracy that are highly unlikely in artificially generated random motif models. This binding preference towards low complexity sequences fits to the previous observation that bipartite motifs tend to bind multiple occurrences of the same sequence.

### Including all binding configurations and cooperativity enhances the accuracy of RBP binding predictions

To assess the value of cooperativity and multivalency, we compared BMF to a 6-mer motif model which scores the sequences by finding the best binding site (Figure S5). Interestingly, for all RBPs but particularly for those that show bipartite binding, BMF’s performance is superior to that of the 6-mer enrichment model. This highlights the value of two distinct BMF features: considering all binding configurations, and including the cooperative effect of multi-domain binding.

### In-vitro bipartite models learned by EMF can predict in-vivo binding

Experimental techniques for measuring RNA binding have individual biases that can be learned by motif discovery tools. This is particularly problematic when evaluating computational methods with many model parameters that can capture complex structures in their input datasets (8). Cross-platform validation, i.e. using binding models trained on an experimental dataset to predicting binding sites in another experimental platform ensures a fair assessment of the quality of motif models. We therefore trained models on HTR-SELEX data to predict binding sites on sequences derived from PARCLIP and eCLIP experiments (40, 24). We compared the performance of BMF to iDeepE (28) and Graphprot (23) (Figure 3A-C). iDeepE is a deep learning tool and Graphprot is based on support vector machines. Thanks their more complex architecture and higher number of parameters, both models are able to learn more complex aspects of the training data, while Graphprot additionally takes the RNA structure as an input. Interestingly, despite these advantages, BMF showed a competitive prediction quality as measured by the area under the receiver operating characteristic curve (AUROC), with a better median AUROC than iDeepE and GraphProt. Simlar results are obtained when replacing AUROC with the area under the precision recall curve (AURPC, Figure S6).

**Figure 3:**
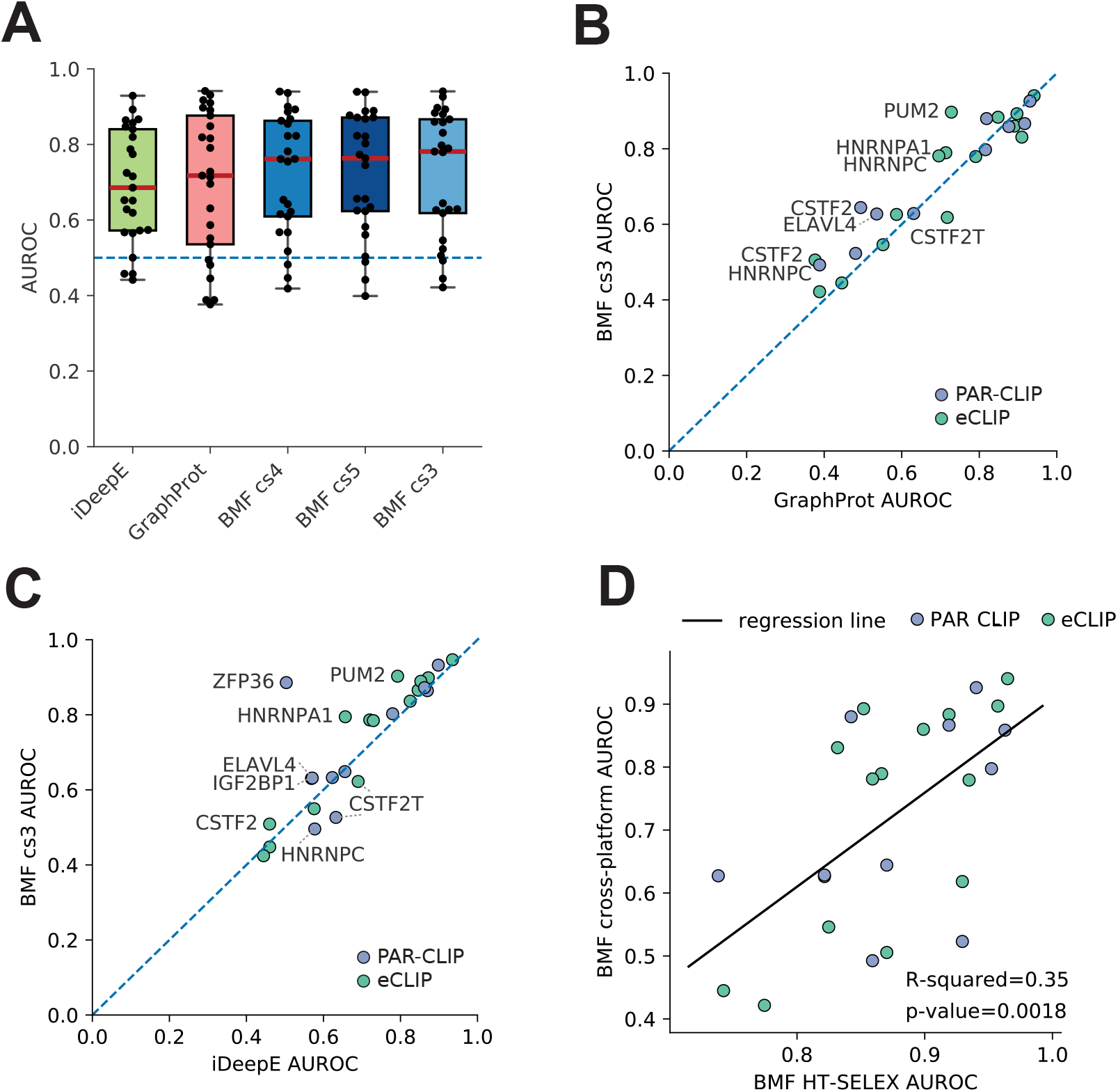
Cross-platform validation shows *in-vitro* BMF motifs can predict *in-vivo* binding sites in transcriptomes. We used BMF, iDeepE, and GraphProt to identify eCLIP and PAR-CLIP RBP binding sites after training their motif models on HTR-SELEX datasets. **(A)** AUROC distribution for iDeepE, GraphProt and BMF with motif sizes ranging from 3 to 5. The tools are sorted based on their median AUROC performance. The values for each RBP dataset is shown with a black dot. **(B)** and **(C)** AUROC from BMF (core size 3) compared to iDeepE and GraphProt. **(D)** BMF AUROC values from cross-validated HTR-SELEX analysis can predict cross-platform performance. Both BMF models are built with core size 3. Linear regression line is marked with black. In all plots AUROC values are averaged over all replicate combinations wherever replicates were available.

Interestingly, generally performance of BMF is best for *k* = 3, although it changes little between core size of *k* = 3, 4 or 5 (Figure 3A, Figure S7). For some RBPs increasing the core size reduced the predictive power for the resulting models. This could be due to over-fitting on biases of the HTR-SELEX data and might be a reason for why the more highly parameterized RNA motif models of GraphProt and iDeepE often do not perform as well as the simpler ones of BMF. On the other hand, longer BMF models, as well as iDeepE and GraphProt, could better learn binding preferences for factors such as CSTF2T that bind more complex RNA sequences. To summarize, BMF can capture RBP specificities with reduced risk of overfitting.

To see whether the core spacing of HTR-SELEX motif models exist in *in-vivo* data, we trained BMF models on the CLIP data and compared them to their *in-vitro* counterparts. Interestingly for the models that were learned well on the HTR-SELEX data (tool-averaged AUROC ≥ 0.8), both the motif core sequences and their distance distribution match between the two experimental platforms (Figure 4). The sequence and\or preferences do vary for other factors with lower AUROC values (Figure S8,S9).

**Figure 4:**
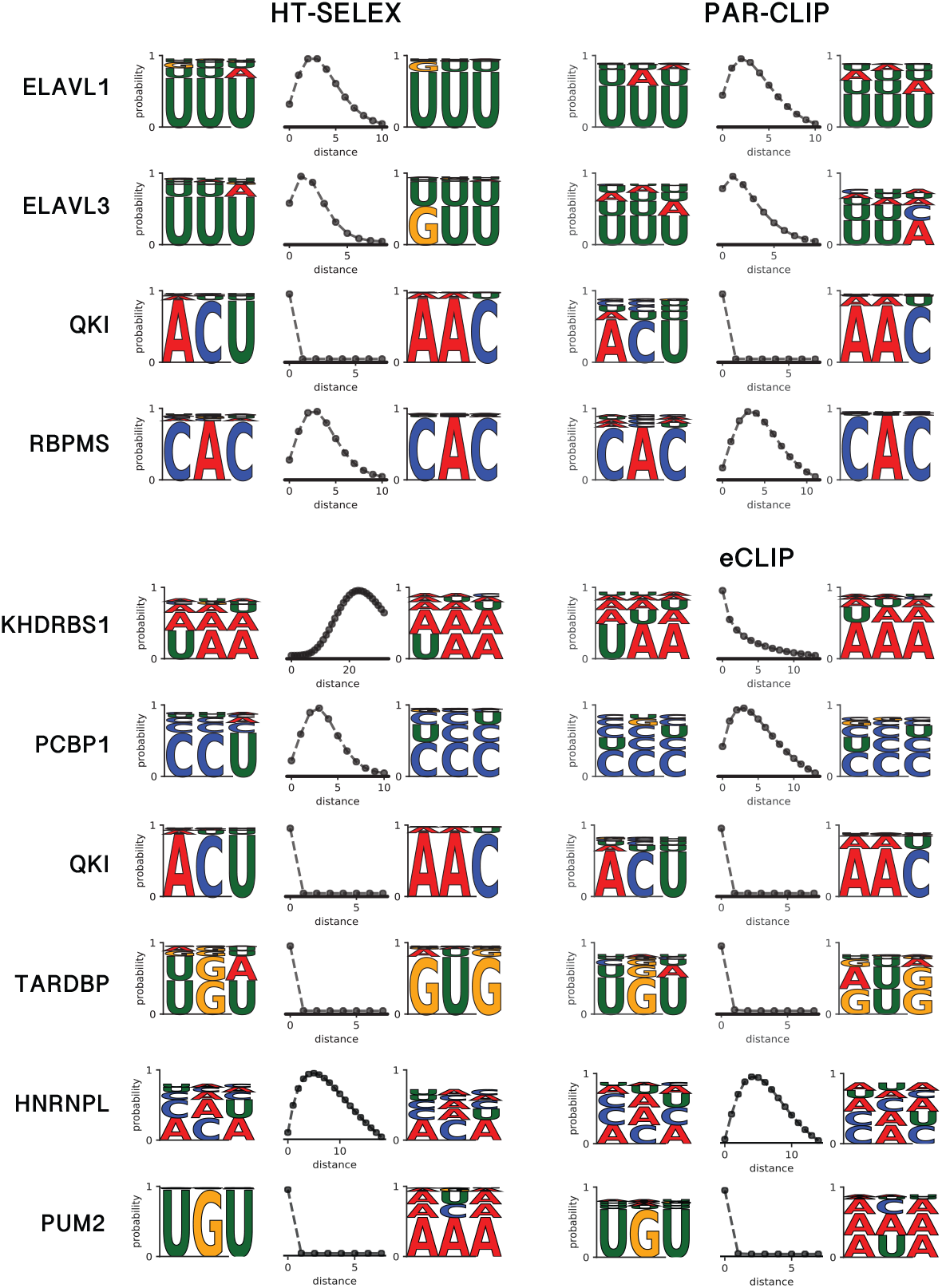
Bipartite motif models learned on *in-vitro* data match their *in-vivo* counterparts. Bipartite motifs are shown for those RBPs in Figure 3 whose best replicate has a tool-averaged AUROC of at least 0.8. The models learned in-vitro and in-vivo match not only in the sequence preference but also the relative positioning of the two motif cores, with the exception of KHDRBS1, which shows a bipartite motif only in the HTR-SELEX data.

A comparison of the AUROC values from the cross-validated HTR-SELEX data (Figure S5) and those from the cross-platform validation shows a correlation between BMF motif quality and its performance in the cross-platform benchmark (p-value=0.0018, Figure 3D). It could help explain why some HTR-SELEX models fail at predicting binding to new sequences, possibly as they have little sequence preference for their target RNA or due to the absence of this information in the HTR-SELEX data. Overall, this shows that BMF can be used to learn RNA motifs from *in-vitro* data to predict binding sites of the protein in the cell despite numerous factors counfounding binding *in vivo*.

## Discussion

We present BMF, the first bipartite motif model to describe multivalent binding preferences in RBPs. The motif models learned on *in-vivo* and *in-vitro* datasets imply the following multipartite binding strategy is common – adapted by about half of RBPs in our datasets – to bind their target RNA molecules: First, these RBPs bind multiple short (3 – 4 nt) RNA segments simultaneously and cooperatively with their multiple RBDs, which can be either on a single chain or part of dimer or oligomer complexes (21, 42). Second, the recognition motifs of their single RNA-binding domains are usually similar (Figure 2). These two aspects make it simple to evolve the sequence features in the target RNAs required for highly specific cooperative binding: a sufficient density of the simple core recognition motifs. We have recently shown that the RBP binding affinity through cooperative binding of multiple RNA-binding domains depends on the motif density on the target RNA with a Hill-like coefficient that is similar in size to the number of binding domains (37, Fig. 4D). Via di- and oligomerization of RBPs the number of cooperatively binding domains and thereby the Hill-like coefficient can be further increased, by which it is possible to distinguish between targets with, say, a core binding motif every 20 versus every 30 nucleotides (e.g 33, Fig. 3EF). An encoding of binding affinity via the density of motifs makes sense for the many RNA-binding proteins for which the precise binding sites on their target RNAs is not important to perform their function.

Mono- and dinucleotide repeats are particularly attractive as target motifs because they possess one binding site per position and per two positions, respectively. The high density of motifs gives rise to high affinities through the combinatorially many possible binding configurations of two or more RNA-binding domains. BMF takes full account of this combinatorial complexity.

A limitation of the evolutionary strategy to bind low-complexity sequences using multiple domains with near-identical motifs is the much smaller number of motifs than can be distinguished, only 64 for length-3 cores. This low number might be sufficient, however, for targeting such RBPs to their RNA targets because specificity is enhanced by compartmentalization – an RBP occurring only in the nucleus cannot bind to cytosolic mRNAs, for example. Furthermore, only a fraction of RBPs is expressed in any one cell type at any one time, in a similar way as the many transcription factors having the same binding affinities are usually expressed in different cells or at different times.

Our results agree with previous studies that reported bipartite motifs in HTR-SELEX and RBNS datasets by counting spaced k-mers of various linker lengths (13, 6). The motifs we report are congruent with those reported before and additionally provide a distance distribution to describe the best binding geometry. The observation that motifs are repetitive and degenerate is also consistent with previous high-throughput studies (6).

Interestingly, BMF motifs were shorter and less complex than those reported by 13. For RBPs for which Jolma et al obtained long motifs (i.e. PCBP1, PUM1, and TARDBP), longer motif cores than 3 nucleotides in BMF could not improve prediction performance in the cross-platform benchmark. This indicates that 3-6 base long motifs would suffice in explaining the sequence specificities for the majority of RBPs.

BMF does not take RNA secondary structure into account. RNA molecules can fold onto themselves and take various tertiary structures (41). It has been shown that some RBPs at least partially identify their target RNA molecules through binding specific structural elements (14, 22). This could further narrow the search space of proteins to fewer potential binding partners and open new ways for cellular regulation. Despite ignoring structure, BMF’s performance is comparable if not better than GraphProt, a tool that includes detailed modelling of secondary structure. We expect that expanding our bipartite motif model to include RNA structure could further improve its predictive power.

Overall, BMF’s performance is promising in the following regards: Owing to its multi-domain binding model BMF can (1) find pairs of sequence motifs over-represented in a sequence set, and can (2) learn the distance between the motif pairs, reflecting the best binding configurations. This information can be further used to (3) asses whether or not an RBP displays bipartite binding. We believe that looking at RNA motifs as combinations of individual low affinity interactions can improve our understanding of RNA regulation in the cell and shed a new light on how some RBPs can find their targets despite the weak sequence and structural preferences of individual domains.

## Code and data availability

The HTR-SELEX data of 13 were downloaded from the European Nucleotide Archive under accession PRJEB25907 (https://www.ebi.ac.uk/ena/browser). The preprocssed eCLIP datasets were collected from the ENCODE at https://www.encodeproject.org (40). PAR-CLIP peaks were obtained from https://github.com/BIMSBbioinfo/RCAS_meta-analysis (24). BMF source code, documentation, and motif models can be found at https://github.com/soedinglab/bipartite_motif_finder.

## Acknowledgements

We thank Christian Roth for his help on AVX-optimization, web server implementation and for discussions on tool development and benchmark design. SSJ acknowledges support from the International Research School for Molecular Biology (IMPRS-MolBio). We acknowledge support by the focus program SPP2191 of the Deutsche Forschungsgemeinschaft.

