## Supplemental Information for "Thermodynamic modeling reveals widespread multivalent binding by RNA-binding proteins"

### Supplementary Methods

#### Parameter Optimization

We learn BMF model parameters by maximizing the likelihood function (equation 9 in manuscript). For an efficient optimization using stochastic gradient descent, we need to be able to compute the partial derivative of the likelihood with respect to model parameters ($\theta$):

$$\frac{\partial LL(\mathbf{\Theta})}{\partial \theta} = \sum_{\mathbf{x} \in \mathcal{X}^+} \frac{1}{Z(\mathbf{x})(1 - Z(\mathbf{x}))} \frac{\partial Z(\mathbf{x})}{\partial \theta} - N^+ \frac{\sum_{\mathbf{x}' \in \mathcal{X}^{\mathrm{bg}}} p_{\mathrm{bg}}(\mathbf{x}') Z(\mathbf{x}')^{-2} \times \frac{\partial Z(\mathbf{x}')}{\partial \theta}}{\sum_{\mathbf{x}' \in \mathcal{X}^{\mathrm{bg}}} p_{\mathrm{bg}}(\mathbf{x}')(1 - 1/Z(\mathbf{x}'))}. \tag{1}$$

We can compute the partial derivatives $\partial Z(\mathbf{x})/\partial \theta$ from the partial derivatives $\partial Z_{\mathrm{A}}(i)/\partial \theta$ and $\partial Z_{\mathrm{A}}(L)/\partial \theta$ according to equation 6 in the manuscript:

$$\frac{\partial Z(\mathbf{x})}{\partial \theta} = \frac{\partial Z_{\mathrm{B}}(x, L-1)}{\partial \theta} + \sum_{i=0}^{L-1} \frac{\partial Z_{\mathrm{A}}(x, i)}{\partial \theta} . \tag{2}$$

BMF parameters ($\theta$) include binding energies of each domain to various $k$-mers as well as concentration parameters ($S$, $r$ and $p$).

In the following, We define $\theta_{k,d}$ as the binding energy of domain $d$ at $k$-mer $k$. Log-likelihood derivatives with respect to binding energies can be computed iteratively by applying the partial derivative operator on the forward algorithm of the dynamic programming (equations 1 and 2 in the manuscript):

$$\frac{\partial Z_{\mathrm{A}}(i)}{\partial \theta_{k,d}} = c_{\mathrm{AB}} e^{-E_A(i)} \left[ \frac{\partial Z_{\mathrm{B}}(i-k)}{\partial \theta_{k,d}} + \sum_{j=0}^{i-k} \frac{\partial Z_{\mathrm{A}}(j)}{\partial \theta_{k,d}} - \left( Z_{\mathrm{B}}(i-k) + \sum_{j=0}^{i-k} Z_{\mathrm{A}}(j) \right) \frac{\partial E_A(i)}{\partial \theta_{k,d}} \right], \tag{3}$$

where

$$\frac{\partial E_A(i)}{\partial \theta_{k,d}} = \delta_{x(i),k}\, \delta_{d,A}\ , \tag{4}$$

and $\delta_{i,j}$ is the Kronecker delta of $i$ and $j$. Similarly we can get the derivatives with respect to $Z_{\mathrm{B}}$:

$$\frac{\partial Z_{\mathrm{B}}(i)}{\partial \theta_{k,d}} = \frac{\partial Z_{\mathrm{B}}(i-1)}{\partial \theta_{k,d}} + \sum_{j=0}^{i-k} c_{\mathrm{B}}(i-k-j)\, e^{-E_B(i)} \left( \frac{\partial Z_{\mathrm{A}}(j)}{\partial \theta_{k,d}} - Z_{\mathrm{A}}(j) \frac{\partial E_B(i)}{\partial \theta_{k,d}} \right)$$
$$+\, c_{\mathrm{AB}}\, e^{-E_B(i)} \left( \frac{\partial Z_{\mathrm{B}}(i-k)}{\partial \theta_{k,d}} - Z_{\mathrm{B}}(i-k) \frac{\partial E_B(i)}{\partial \theta_{k,d}} \right), \tag{5}$$

where

$$\frac{\partial E_B(i)}{\partial \theta_{k,d}} = \delta_{x(i),k}\, \delta_{d,B}\ . \tag{6}$$

These derivatives are computed iteratively via dynamic programming similar to $Z_{\mathrm{A}}$ and $Z_{\mathrm{B}}$. They are initialized to zero for:

$$\frac{\partial Z_{\mathrm{A}}(i)}{\partial \theta_{k,d}} = 0 \text{ for all } i \in \{0, ..., k-2\} \tag{7}$$
$$\frac{\partial Z_{\mathrm{B}}(i)}{\partial \theta_{k,d}} = 0 \text{ for all } i \in \{0, ..., k-2\}\,. \tag{8}$$

Similarly, we can derive the partial derivative in respect to the concentration parameters ($\theta_c$):

$$\frac{\partial Z_{\mathrm{A}}(i)}{\partial \theta_c} = c_{\mathrm{AB}}\, e^{-E_A(i)} \left( \frac{\partial Z_{\mathrm{B}}(i-k)}{\partial \theta_c} + \sum_{j=0}^{i-k} \frac{\partial Z_{\mathrm{A}}(j)}{\partial \theta_c} \right)\ , \tag{9}$$
$$\frac{\partial Z_{\mathrm{B}}(i)}{\partial \theta_c} = \frac{\partial Z_{\mathrm{B}}(i-1)}{\partial \theta_c} + \sum_{j=0}^{i-k} e^{-E_B(i)} \left( \frac{\partial Z_{\mathrm{A}}(j)}{\partial \theta_c} c_{\mathrm{B}}(i-k-j) + Z_{\mathrm{A}}(j) \frac{\partial c_{\mathrm{B}}(i-k-j)}{\partial \theta_c} \right)$$
$$+ \frac{\partial Z_{\mathrm{B}}(i-k)}{\partial \theta_c} c_{\mathrm{AB}} + Z_{\mathrm{B}}(i-k) \frac{\partial c_{\mathrm{AB}}}{\partial \theta_c}\ . \tag{10}$$

The partial derivative $\partial c_{\mathrm{B}} / \partial \theta_c$ with respect to concentration parameters $S$, $r$ and $p$ are (according to equation 5 in the manuscript):

$$\frac{\partial c_{\mathrm{B}}(d)}{\partial S} = \frac{\Gamma(d+r)}{\Gamma(d+1)\Gamma(r)} p^r (1-p)^d, \tag{11}$$
$$\frac{\partial c_{\mathrm{B}}(d)}{\partial p} = S \times \left(\frac{r}{p} - \frac{d}{1-p}\right) \times \exp\big( \log \Gamma(d+r)$$
$$- \log \Gamma(d+1) - \log \Gamma(r) + d \log(1-p) + r \log p \big), \tag{12}$$
$$\frac{\partial c_{\mathrm{B}}(d)}{\partial r} = S \times \big( \psi(d+r) + \log p - \psi(r) \big) \times \exp\big( \log \Gamma(d+r)$$
$$- \log \Gamma(d+1) - \log \Gamma(r) + d \log(1-p) + r \log p \big), \tag{13}$$

where $\Gamma$ is the gamma function and $\psi$ is its logarithmic derivative, also known as the digamma function. Note that for numerical accuracy, we have calculated the derivatives of the $\exp(\log c_{\mathrm{B}}(d))$ function, with respect to $p$ and $r$.

Overall, these equations allow us to iteratively compute for any sequence $\mathbf{x}$ the partial derivatives $\partial Z_{\mathrm{A}}(i)/\partial\theta$ and $\partial Z_{\mathrm{B}}(L)/\partial\theta$ with respect to all parameters and hence derivatives of the partition function $Z(x)$ and those of the likelihood.

The thermodynamic model contains a simplification. We assume that one of the domains (A) always binds upstream of the other domain (B) and that the binding configurations A-B and B-A do not *both* contribute appreciably to the binding probability. This seems like a very plausible assumption considering that the linkers between structural domains are usually quite short, and changing the order of binding would usually result in an impossible or much less favorable (tighter) configuration of the RNA chain.

### Calculation of motif entropy

To derive the entropy for each bipartite motif model, we calculate the weighted probability for each base as

$$P_b = \frac{\sum\limits_{x\in k\text{-mers}} n_{b,x} p_x + \sum\limits_{y\in k\text{-mers}} n_{b,y} p_y}{\sum\limits_{b\in N} \Big( \sum\limits_{x\in k\text{-mers}_A} n_{b,x} p_x + \sum\limits_{y\in k\text{-mers}} n_{b,y} p_y \Big)}, \tag{14}$$

where $N$ is the set of nucleotides ($\{A, C, G, U\}$). We calculate the entropy as

$$\text{Entropy} = -\sum_{b\in N} P_b \log_2 P_b \,. \tag{15}$$

To establish a baseline for the observed entropy values, we generated artificial bipartite motifs where the $k$-mer probabilities are taken from the observed probabilities of an experimental set but the $k$-mers were shuffled. We generated 10,000 such motifs and used the resulting entropy distribution as a baseline for motif complexity.

### Calculation of motif repetitiveness

To quantify the degree of sequence repetitiveness in BMF models, we calculate the highest average probability of observing a repetitive 3-mer (i.e. 'AUA', 'UUU', or 'CGC') as

$$R = \max_{a,b\in N} \left( \sqrt{p_A(aba) + p_A(bab))(p_B(aba) + p_B(bab)} \right), \tag{16}$$

where $N$ is the set of nucleotides ($\{A, C, G, U\}$), and $p_A$ and $p_B$ are BMF probabilities for the first and second motif core respectively. To establish a baseline for the observed repetitiveness values, we calculated this metric for 10,000 artificial bipartite motifs, generated as described above.

### BMF comparison with single-occurrence motif model

To estimate the effect of considering all binding configurations and including cooperativity in BMF, we compared its cross-validated classification performance with a 6-mer motif model. We created training and test sets by splitting the HTR-SELEX data with an 80 to 20 ratio. For the 6-mer model, we calculated enrichment factors for each 6-mer in training data and scored the test sequences by the most enriched 6-mer motif. Similarly, we trained BMF with core size 3 on the training data and used the learned models to predict the binding scores for each sequence in the test set. To compare each model's classification power, we calculated the area under the receiver operating characteristic curve (AUROC) for all RBPs in the dataset.

### Supplementary figures

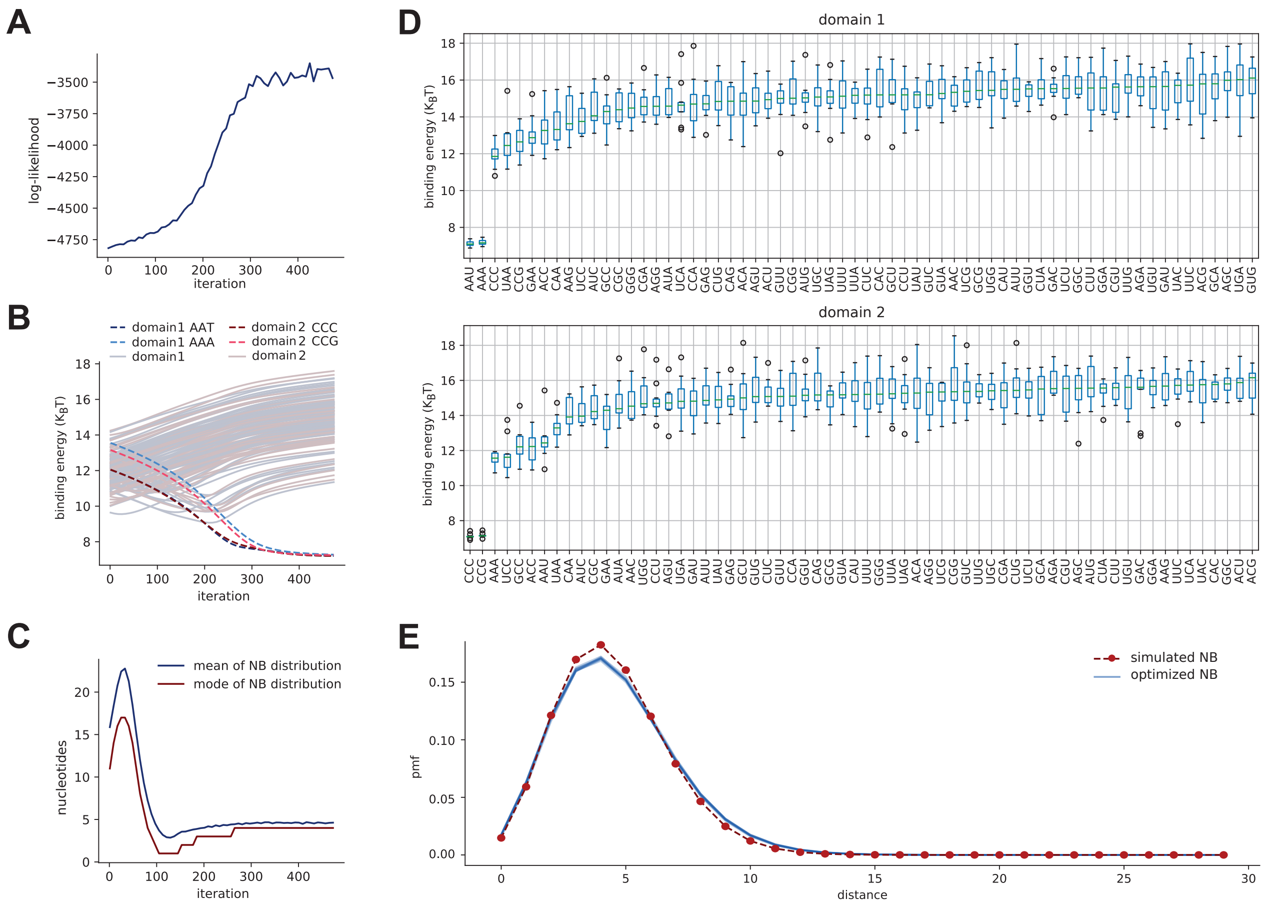


Figure S1: **BMF can reliably learn planted motifs in synthetic data.** We planted AA(A/U) followed by CC(C/G) with a distance distribution around 4 in 2000 randomly generated sequences of length 40. **(A)** Log-likelihood function increases over the iterations until reaching a plateau at the end of optimization. **(B)** The binding energies of all 3-mers are shown over BMF iterations for both binding domains. The 3-mers representing the implanted motifs are shown with brighter blue (first domain) and red (second domain)dash lines. The final values retrieved after optimization is notably lower for the highlighted 3-mers. **(C)** The mean (in blue) and mode (in red) of the NB distribution is shown over BMF's optimization iterations. The correct distance distribution is found when the LL and the energy parameters reach their plateaus. **(D)**, and **E** The distribution of final BMF parameters upon 10 random parameter initializations and subsequent optimization. Regardless of the choice of initial parameter values, BMF ends in the same optimum point in the parameter landscape.

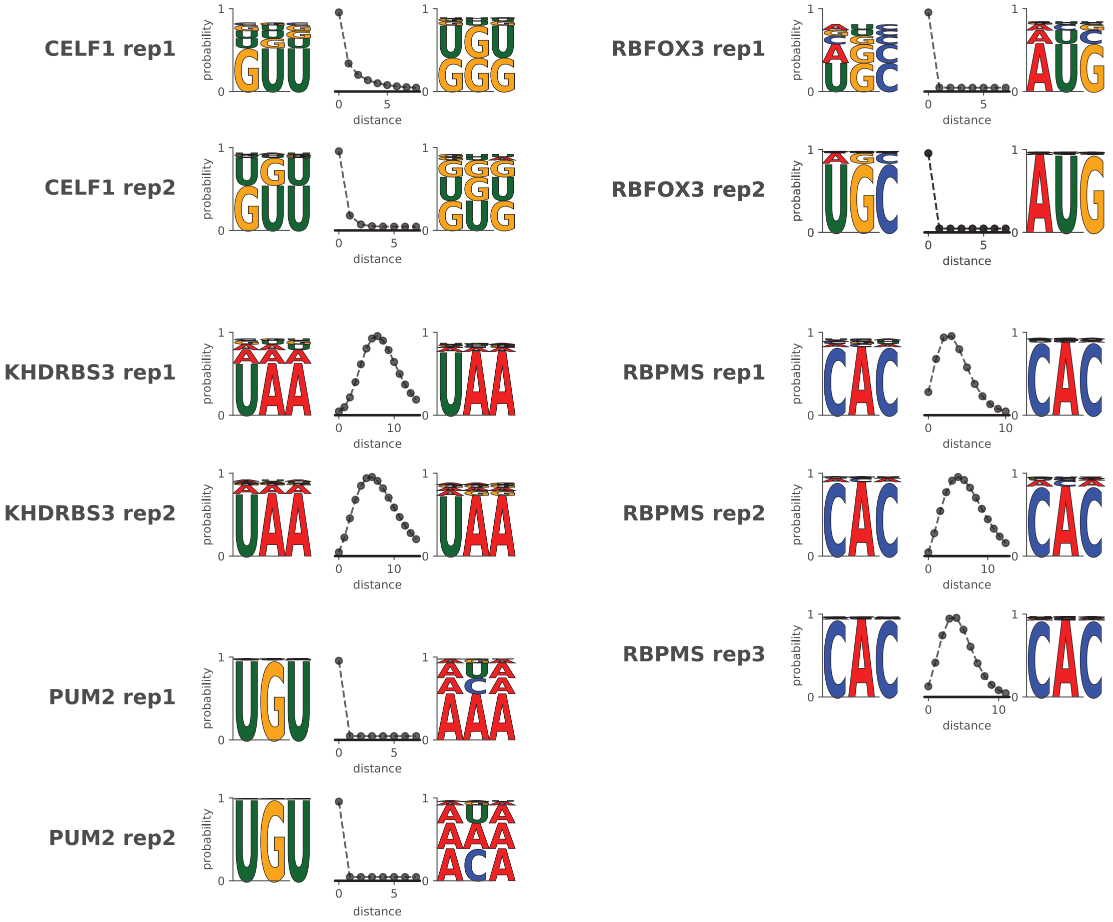


Figure S2: **Experimental HTR-SELEX replicates generate the same bipartite motif models**. Biparite binding models are shown for factors in Figure 2 for which an experimental replicate was available. The models generated for all HTR-SELEX datasets can be found in BMF GitHub repository: `https://github.com/soedinglab/bipartite_motif_finder/blob/main/data/HTRSELEX_motifs.pdf`.

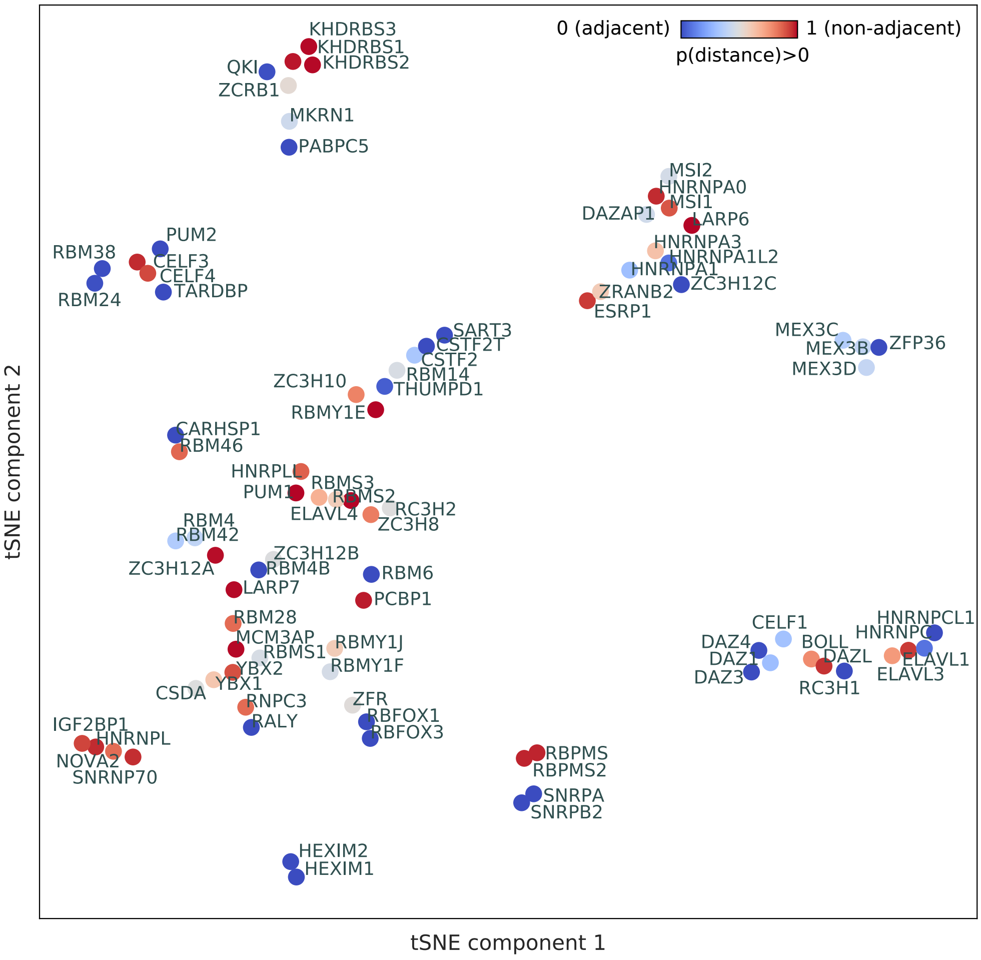


Figure S3: **RBPs in the same family have similar BMF motifs**. RBPs are clusters according to their sequence identity measured as pairwise Pearson correlation between 3-mer probabilities. Two dimensional embedding is generated via tSNE [1]. RBPs are color-coded based on the domain positioning in the NB models, as in Figure 2B, with adjacent cores colored in blue and bipartite motifs in red.

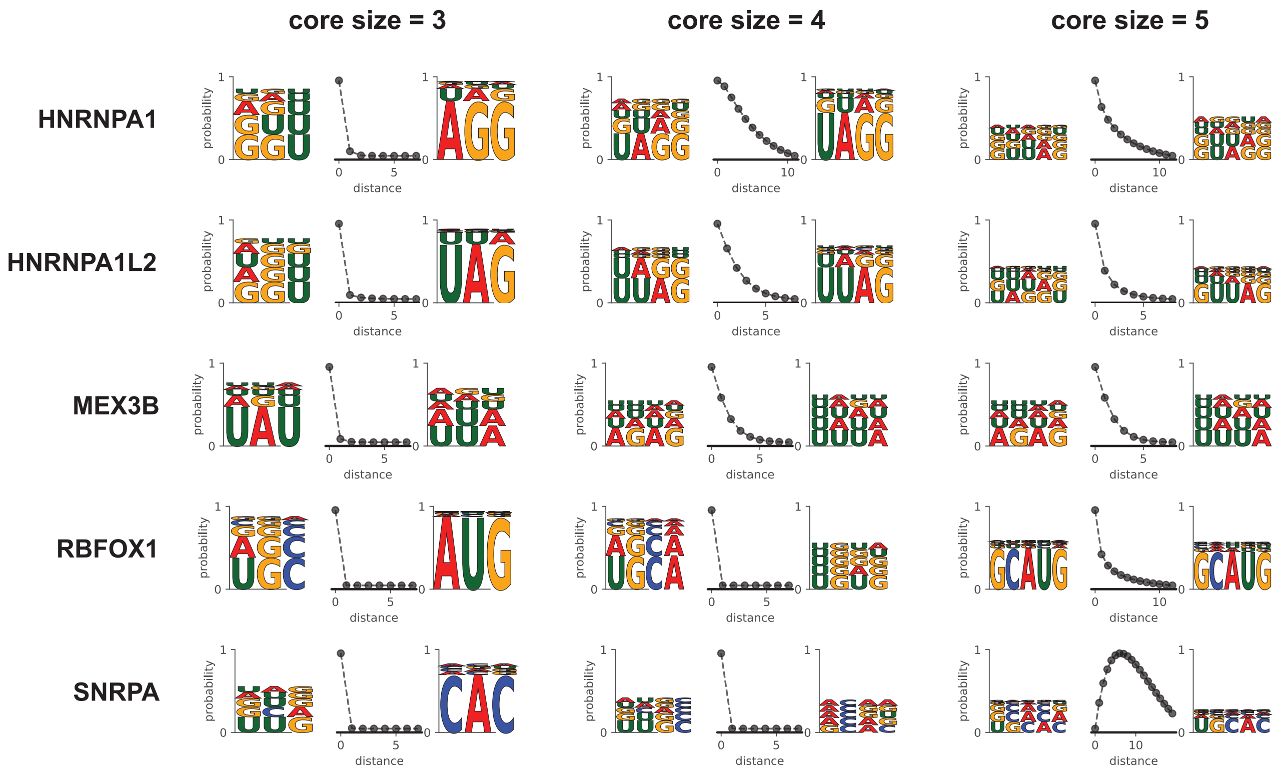


Figure S4: **Bipartite binding behaviour can arise when biulding longer sequence models**. Some RBPs in the HT-SELEX dataset have adjacent cores when building BMF models with 3-mers, but show bipartite binding for 4-mer and\or 5-mer models.

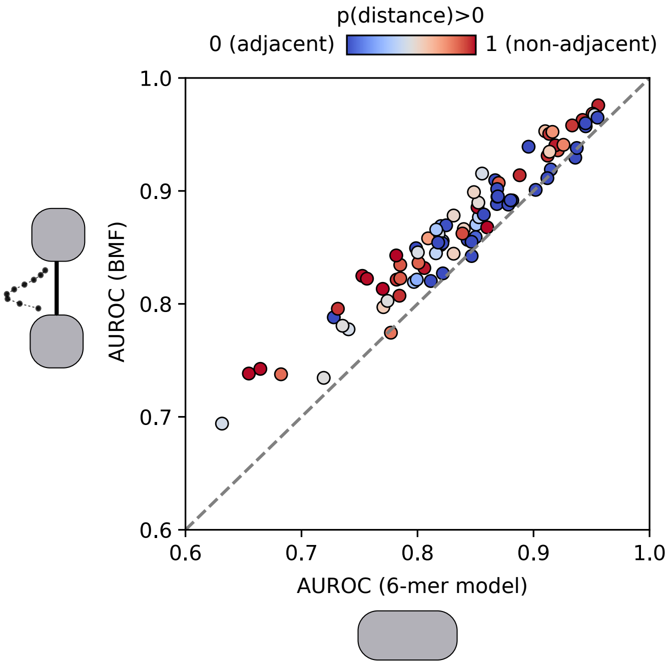


Figure S5: **Incorporating cooperativity and multivalency boosts performance of RBP binding models.** AUROC values are calculated by predicting binding sites in held-out sequences of HTR-SELEX datasets (80%-20% split for training and testing). BMF with core size 3 is compared to a single-occurence per sequence 6-mer model. RBPs are color-coded based on the domain positioning in the NB models, as in Figure 2B, with adjacent cores colored in blue and bipartite motifs in red.

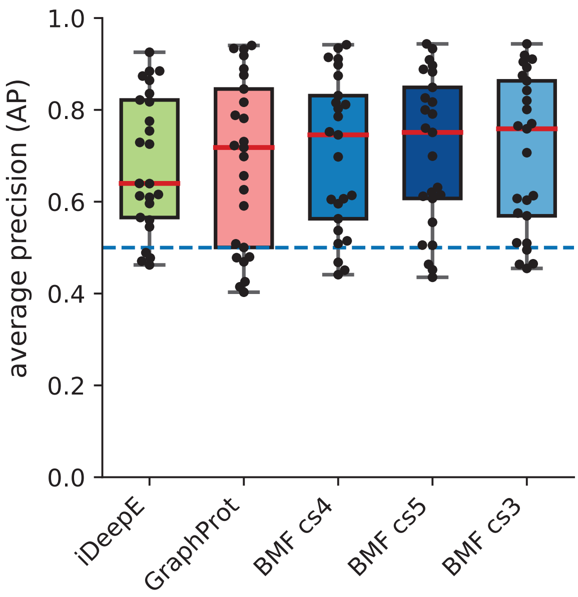


Figure S6: **Average precision (AP) scores for iDeepE, GraphProt and BMF with motif sizes ranging from 3 to 5.** We used BMF, iDeepE, and GraphProt to identify eCLIP and PAR-CLIP RBP binding sites based on the models trained on HTR-SELEX datasets. The tools are sorted based on their median AP scores (red lines). The AP score for each RBP dataset is shown with a black dot.

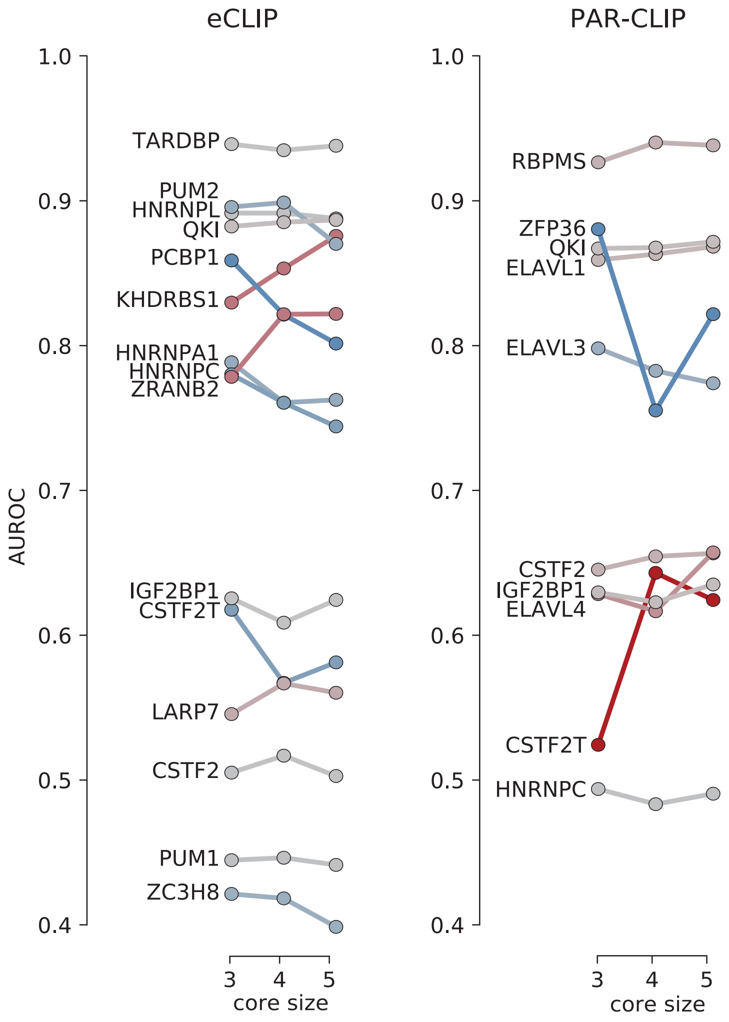


Figure S7: **Comparison of cross-platform AUROC values for BMF models with core sizes 3 to 5.** An increase in AUROC with increasing motif length is marked with red and a decrease with blue.

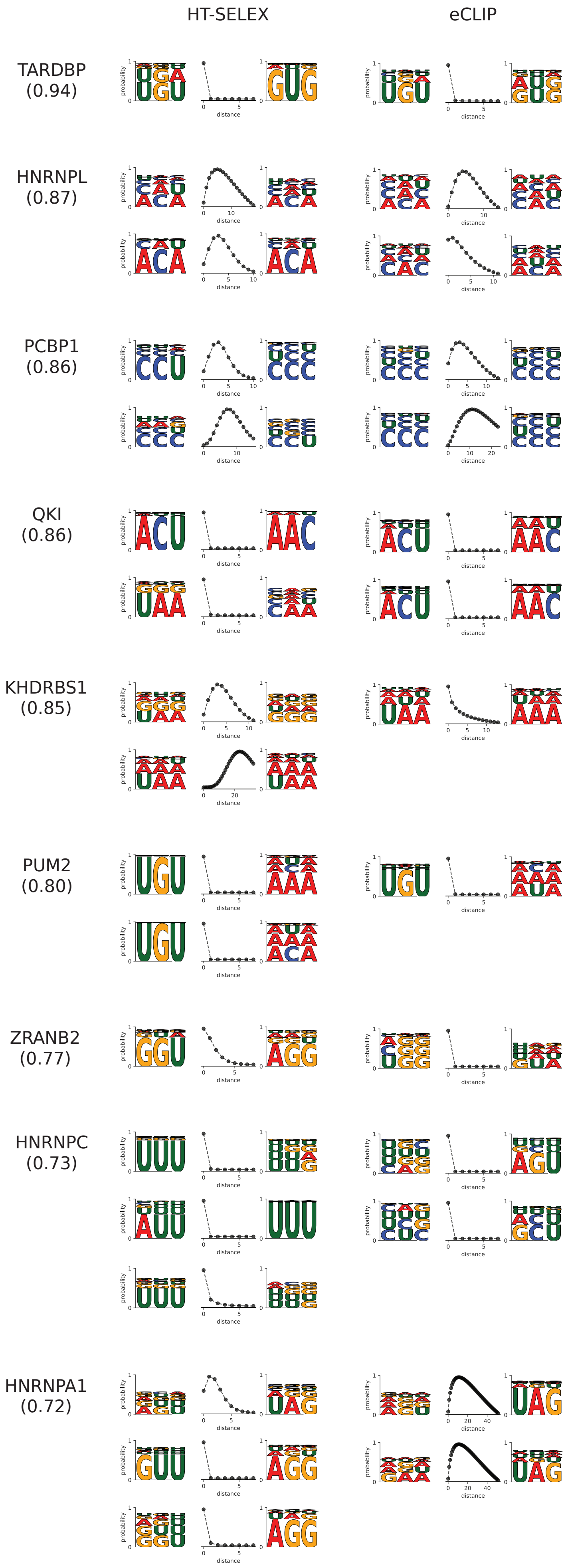
HT-SELEX
eCLIP
TARDBP
(0.94)
HNRNPL
(0.87)
PCBP1
(0.86)
QKI
(0.86)
KHDRBS1
(0.85)
PUM2
(0.80)
ZRANB2
(0.77)
HNRNPC
(0.73)
HNRNPA1
(0.72)
probability
distance

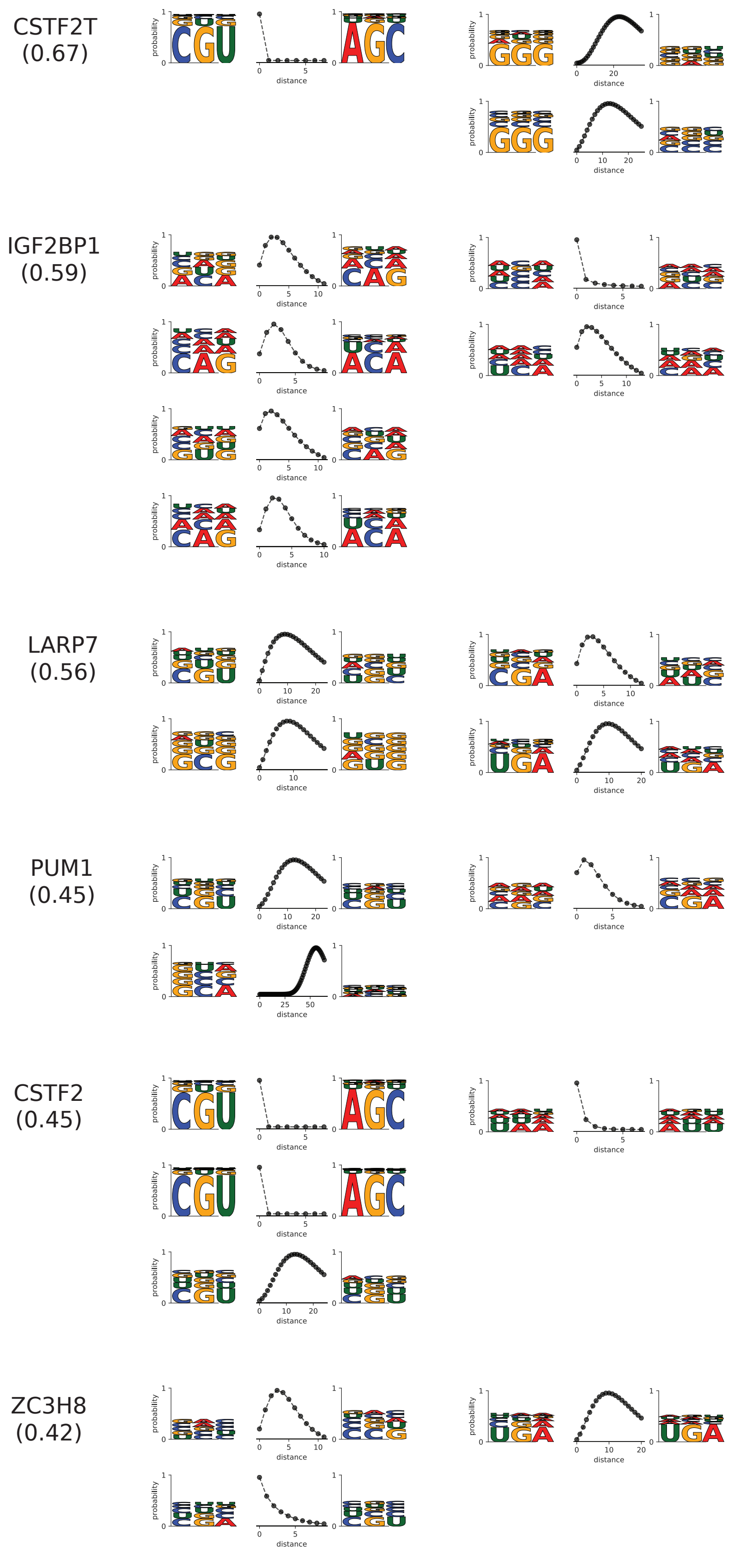


Figure S8: **Comparison of HTR-SELEX and eCLIP BMF logos**. BMF logos are sorted according to their cross-platform AUROC performance (shown in parenthesis), which is an average between BMF (with core size 3), Graphprot, and iDeepE. BMF logos were generated for all available replicates of each experimental technique.

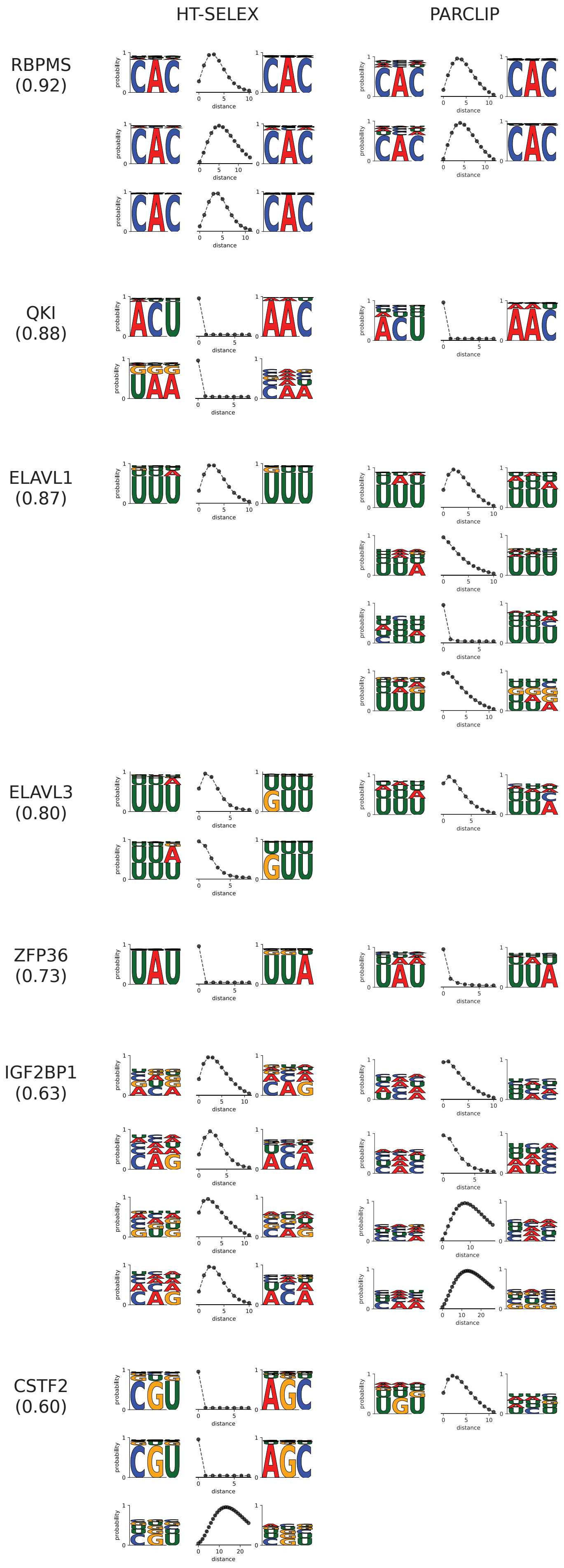
HT-SELEX
PARCLIP
RBPMS
(0.92)
QKI
(0.88)
ELAVL1
(0.87)
ELAVL3
(0.80)
ZFP36
(0.73)
IGF2BP1
(0.63)
CSTF2
(0.60)
probability
distance

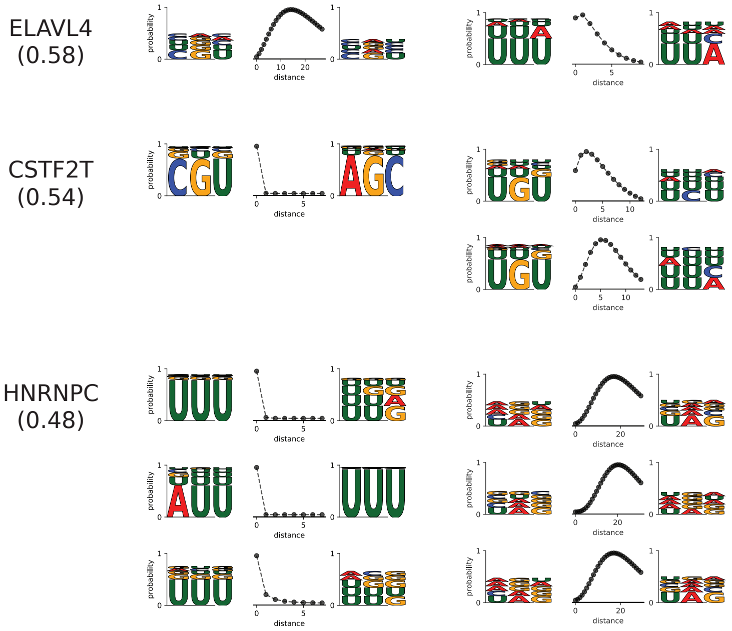


Figure S9: **Comparison of HTR-SELEX and PAR-CLIP BMF logos**. BMF logos are sorted according to their cross-platform AUROC performance (shown in parenthesis), which is an average between BMF (with core size 3), Graphprot, and iDeepE. BMF logos were generated for all available replicates of each experimental technique.
